# Well-ST-seq: cost-effective spatial transcriptomics at cellular level and high RNA capture efficiency

**DOI:** 10.1101/2023.06.28.546974

**Authors:** Nianzuo Yu, Zhengyang Jin, Chongyang Liang, Junhu Zhang, Bai Yang

## Abstract

Spatial transcriptomic technologies are promising tools to reveal fine anatomical profiles of tissues. As for the methodologies based on barcoded probe arrays, improving the balance among probe barcoding complexity and cost, gene capture sensitivity, and spatial resolution can accelerate the spreading of spatial transcriptomic in basic science and clinical work. Here, based on miniaturized microfluidic and microarray technologies, we constructed a spatially cellular-level RNA-capture probe arrays. Owing to the predetermined and cost-effective probe fixation characteristics of the methodology, the consumable cost and fabrication time of the probe array can be reduced to $1.21/mm^2^ and approximately 2 hours, and the preparation process does not rely on large precision instruments. Moreover, the efficiency of the transcript captured by the probe array is even comparable to conventional single-cell RNA sequencing. Based on this technology, we achieved the spatial transcriptome expression mapping and gained insight into spatial cell heterogeneity of the mouse hippocampus.

## Introduction

Defining gene expression patterns from a spatial perspective can provide insights into the development and maintenance of complex tissue architectures and the molecular characterization of pathological states^1,2^. Spatial transcriptomic methodologies can display gene expression in different regions of tissue sections, and reveal activated signaling pathways in fine pathological regions^3,4^. Compared with single-cell RNA sequencing (scRNA-seq) technology, spatial transcriptomics have completed the technological innovation of pathological digitization combined with pathological imaging, playing an important role in the development of diagnostic markers, drug resistance sites, targeted drugs, immunotherapy, and other emerging fields^5-7^.

Early spatial transcriptomic sequencing methods are based on single-molecule fluorescence imaging techniques, such as RNA in situ hybridization, which allows the detection of only one or a handful of target molecular species at a time^8^. The development of the electron microscopy has greatly promoted the progress of the imaging-based spatial omics methods. At present, the spatial techniques, such as FISSEQ and MERFISH6, can even simultaneously detect the spatial distribution of 10,000 RNAs in tissue sections, which greatly deepens our understanding of pathophysiological processes^9,10^. Considering the requirements of the imaging technologies for super-resolution microscopy, SeqFISH+ combined the hydrogel embedding and spatial barcoding strategies, and reduced the imaging requirements to the level of standard confocal microscopes, improving the application scope of the imaging-based spatial sequencing techniques^11^.

Spatially patterned DNA barcoding technology provides an innovative detection platform for spatial transcriptomics^12^. Compared with imaging-based sequencing technologies, the information extraction of transcriptomes in the strategy relies on *in-situ* capture of tissue RNA, which does not require multiple rounds of fluorescence imaging processes in tissues, and is simpler in operation and easier to train^13^. The method is also compatible with Next-generation sequencing (NGS) sequencing, and the detection results are unbiased, which is therefore well suited for exploring a new system. In recent years, several remarkable methodologies, such as Slides-seq^14^, HDST^15^, Seq-Scope^16^, Stereo-seq^17^, have improved the spatial resolution of the DNA spot to single-cell or sub-cell level, while the high-resolution NGS-based methods still require cost-limiting fluorescence imaging process to decode the barcoded sequences on the spots. Pixel-seq uses a stamp method to copy the decoded RNA-capture sequences to another hydrogel surface, reducing the frequency of the fluorescence decoding procedure, while the repeated stamping would lose ∼15% of features after 50 cycles^18^. DBiT-seq directly marks the position of RNA in tissue sections, which omits the fluorescence decoding process^19^. So far, the cost and operational versatility are still the main factors restricting the further promotion of the NGS-based spatial transcriptomic technology. The current methods either require large precision instruments to spot or decode patterned RNA-capture sequences, or need to be familiar with microfluidic manipulation after tissue sectioning, and cannot simultaneously take into account the low-cost preparation of barcode arrays and the straightforward *in-situ* RNA capture operations^20^.

Here, the microwell arrays filled with hydrogel beads are used to connect barcoded RNA-capture probes and realize low-cost spatial transcriptomics at cellular level, designated as Well-ST-seq. The barcoded sequences of the hydrogel beads are encoded by the microfluidic technology, avoiding the use of large-scale instruments, such as fluorescence microscopy or point printer, and the sequencing procedure is compatible with *in-situ* RNA capture operation and NGS technology. Notably, the network of hydrogel beads can be grafted with high-density bioactive carboxyl groups, and the output of the captured transcriptome can reach 3896 UMIs in 10 μm resolution spot, which is outstanding among the reported spatial methodologies. When aggregated into single-cell areas, the detected transcriptome of Well-ST-seq (∼5100 UMIs per cell on average) is comparable to conventional scRNA-seq technology.

## Result

### Well-ST technology overview

Well-ST-seq workflow is mainly divided into three steps (Fig.1A): beads arrangement, microfluidic encoding, and tissue section sequencing. The beads arrangement is to uniformly disperse the beads into microwell arrays, and the microfluidic encoding is to connect spatially barcoded RNA-capture molecules on the surface of the beads.

**Fig.1.**
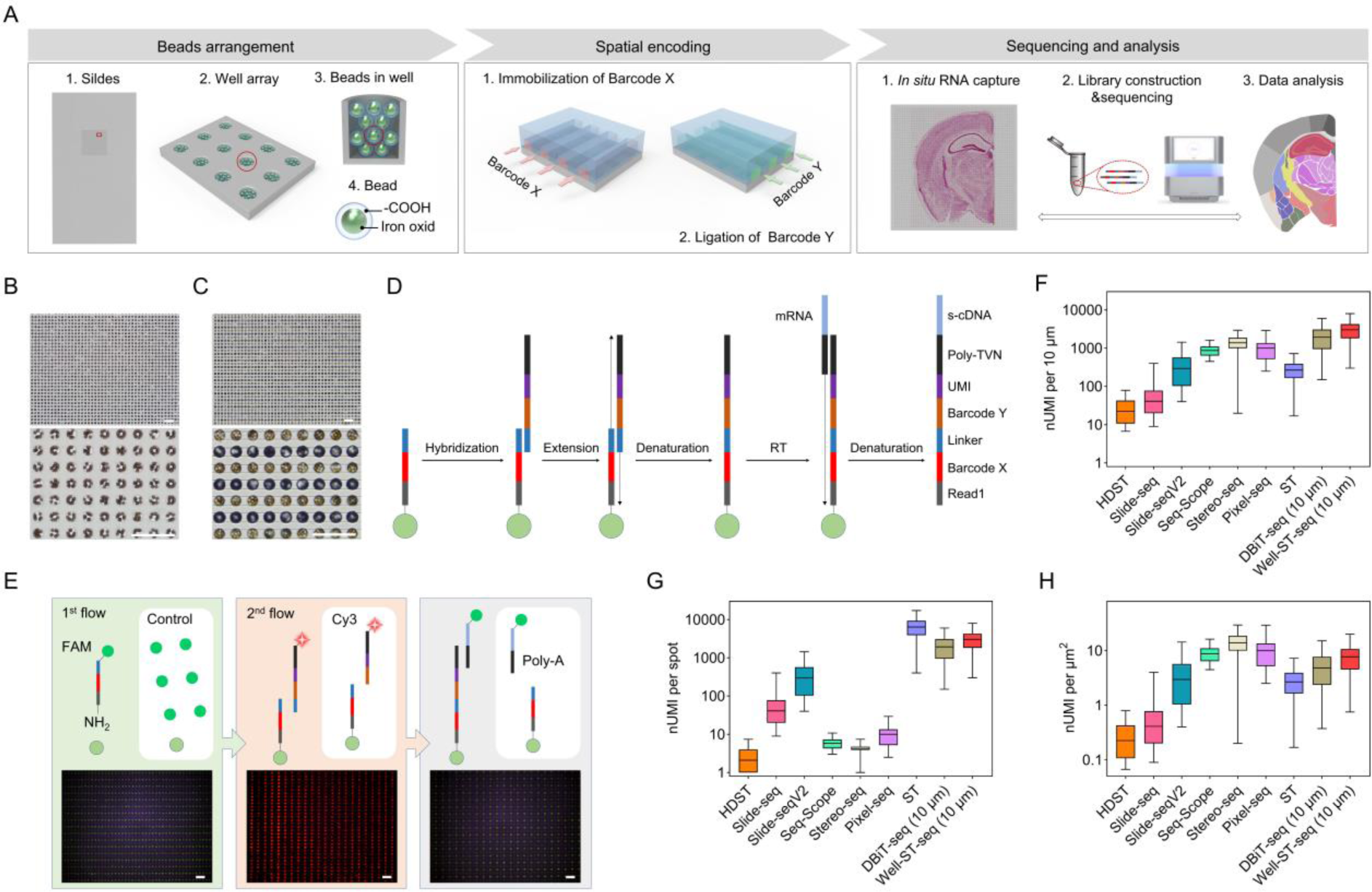
Design and validation of Well-ST-seq. A, Bead arrangement. The particle solution was loaded onto to the surface of the hydrophilic microwell array. After the liquid was volatilized, a flat PDMS sheet was used to remove the beads outside the microwell array, and then the beads array was obtained. The bead is a paramagnetic particle, and the hydrogel network with high-density carboxyl groups is grafted on the surface of the particle. Spatial encoding. The microfluidic chip with 60 parallel microchannels was aligned on the microwell array, and a set of barcode X solutions were introduced into the parallel microchannels, respectively. After the fixation of barcode X, the microchannel was replaced by another microchannel to introduce a second of barcode Y along the orthogonal direction. After the extension and denaturation of barcode X and barcode Y, spatially barcoded RNA-capture spots were obtained. Sequencing and analysis. Frozen tissue section is loaded onto the surface of the bead array, followed by fixation, permeabilization for *in-situ* RNA capture, reverse transcription, and NGS sequencing. Finally, the spatial transcription information of tissue sections is presented through data analysis. B, Dispersed beads in the microwell array. C, The microfluidic chip with parallel microchannels array aligned on the microwell array, and the fluidity and independence of the microchannel array were verified by alternately introducing fluid into the microfluidic chip. D, The sequence composition changes of the RNA-capture probe and the RT process of the transcripts. E, Validation of spatial barcoding for the 10 μm-resolution microwell array using fluorescent probes. The fluorophore-labeled barcodes and the control solutions were alternately introduced into the microchannel array, and only the region where the barcode flowed through showed fluorescence signal. F-G, UMI count distribution. Well-ST-seq is compared to reported HDST, Slide-seq, Slide-seqV2, Seq-Scope, stereo-seq, Pixel-seq, Visium, and DBiT-seq (10 μm) datasets with different spot/pixel sizes.

The beads arrangement starts from the fabrication of microwell arrays. The arrays can be designed as 60×60 independent spots, and its spatial resolution can be set to 10-50 μm as required. The height of the microwell and the diameter of the hydrogel beads can be selected to 15-20 μm and 3-5 μm, respectively, and the microwell with 10 μm diameter can accommodate 10-15 beads (Fig.1B). Compared with a single large-sized bead, multiple small-sized beads can hold larger specific surface area, which improved the modification density of RNA-capture probes. To further increase the quantity of the probes, the bioactive groups for fixing the probes were designed on the branched chains of the repeat units of the hydrogel networks on beads.

The spatial encoding of the patterned RNA-capture probes on beads was achieved by microfluidic technology (Fig.1A). The microfluidic chip contained 60 parallel polydimethylsiloxane (PDMS) microchannels (Fig.S1A,C), which were aligned on the microwell array (Fig.1C), and a set of barcode X solutions were introduced into the parallel microchannels, respectively. Each barcode X molecule composes of a bioactive linking group for fixing the barcode X to the hydrogel networks of beads, the PCR adaptor sequence (22-mer), a distinct spatial barcode Xi (i = 1-60, 8-mer), and a ligation linker (10-mer, Fig.1D). After the fixation of the barcode X, the microchannel was replaced by another microchannel to introduce a second set of barcode Y along the orthogonal direction of the barcode X fluids (Fig.S1B,D), each barcode Y molecule containing a ligation linker for hybridizing with barcode X, a distinct spatial barcode Yj (j = 1-60, 8-mer), a unique molecular identifier (UMI), and an oligo-polyA sequence for generating oligo-dT sequence that can capture tissue polyA-tailed RNA (Fig.1D). After the extension and denaturation of the barcode X and barcode Y, spatially barcoded RNA-capture sequences were respectively fixed on the beads in the microwell array, each containing a distinct combined sequence of barcode Xi and Yj (i = 1-60, j = 1-60).

Frozen tissue sections (10 μm thickness) were loaded onto the surface of the beads array, followed by fixation, permeabilization for *in-situ* RNA capture, reverse transcription (RT) plus amplification, and then the cDNA molecules were linked to the corresponding positions of the spatial barcodes. After the collection of the marked cDNA, the library preparation and NGS sequencing can be performed. Finally, the spatial transcription information of tissue sections can be presented through computational analysis.

### Evaluation of the flow barcoding process

Evaluation of the flow barcoding process is mainly based on two indicators: the mobility of fluids in the microchannel and the diffusion of fluids between the microchannels. To prevent fluids from driving the beads to the adjacent microwells or blocking the microchannel, the height of the microchannel was set to be less than the diameter of the beads. In addition, a magnet was placed at the bottom of the microwell slide to further confine the beads in the microwell. The barcode solution can flow freely without obvious fluid leakage between the microchannels (Fig.1C, Fig.S1E,F). In order to further prove the flow independence of the microchannel array, we adopted the fluorescence characterization method with higher precision, and conjugated the barcode Xi (i = 1-60) and RNA standards (oligo-polyA) with fluorophore FAM, and barcode Yi (i = 1-60) with fluorophore Cy3. First, FAM-labeled barcode X and FAM dye solutions were alternately introduced into the microchannel array, and the concentration of the barcode solutions was set to twice the sequence concentration used to accelerate the rate of the molecule diffusion. The fluorescence signal was observed only on the beads where barcode X flows through (Fig.1E). Next, we used a similar method to introduce the Cy3-labeled barcodes Y solution containing and without a linker over the beads array grafted with barcode X. It was found that only the beads region in contact with the linker of the barcode Y showed fluorescence signal. Finally, we successively immobilized barcode X and barcode Y on the beads surface, and found that fluorescence signal only appeared on the beads region where both barcodes X and barcodes Y flowed through, which validated spatially confined delivery and binding of barcodes to beads array using microfluidic technology.

### Quality assessment of Well-ST-seq sequencing data

The PCR amplicons were used to analyze the size distribution of sequence library construction, which showed the peaks around 300-600 bp (Fig.S2A). The NGS technology was used to detect the sequence library, and the reads were routinely filtered to extract barcode X, barcode Y and UMIs for each spot. The processed reads were demultiplexed, trimmed, and mapped against the mouse genome (GRCh38, Gencode M11 annotation) using the pipeline reported previously^21^, and the UMI and detected genes of each spot were counted. In the processed data of the Well-ST-seq experiment at 10 μm resolution (Well-ST-seq-10 μm), we detected an average of 3896 UMIs per spot (Fig.S2B), which was superior to other existing 10 μm-resolution spatial transcriptomic technologies (Fig.1F), including Slide-seq (∼ 60 UMIs)^14^, Slide-seqV2 (550 UMIs)^22^, and DBiT-seq (10 μm resolution, ∼ 3500 UMIs)^19^. Although the number of UMIs measured per spot of the Visium technology was higher than that of the Well-ST-seq^13^ (Fig.1G), the spot size in Visium was larger than 20 times that of the Well-ST-seq-10 μm, and the UMI capture per unit area was much lower than that of the Well-ST-seq-10 μm (Fig.1H). Considering that the sequencing depth of the library sequence was only 30 g, the maximum possible Well-ST-seq capture efficiency should be even higher than the currently presented data. We also showed the saturation curves of Well-ST-seq-10 and Well-ST-seq-20 μm, and found that they were nearly identical, proving the consistency and low variability of the technology (Fig.S2C,D). Compared with the Visium and DBiT methods, the saturation curves showed a similar trend, but Well-ST-seq could detect more genes. Similar to the detection results observed in Visium and DBiT, Well-ST-seq also existed detection bias dependent on the gene length (Fig.S2E).

### Well-ST-seq-20 μm reveals the spatial transcriptomic mapping of a mouse brain

As proof of principle of the technological strength of Well-ST-seq, we presented the spatial transcriptomic map of the mouse brain in a tissue section. Its adjacent section was used for H&E staining (Fig.2A). The spatial map of UMI and gene counts showed an average of 5764 UMIs and 3000 genes per spot, respectively (Fig.S3A). To benchmark the Well-ST-seq data, we examined a set of genes from different regions of mouse brains, including Meis2, Tnnt1, Kcne2, Slc5a7, Spink8, which were expressed in the thalamus (RT), thalamus (TH), fiber tracts (FT), third ventricle (V3), and hippocampal formation (HPF) regions of the mouse brain (Fig.2B,S3B), and we found that the gene distribution presented by Well-ST-seq agreed with the *in-situ* hybridization (ISH) data from the Allen Mouse Brain Atlas (AMBA). Unsupervised clustering of all the spatial spots was characterized by specific marker genes and identified 9 distinct clusters (Fig.2C,D), which corresponding to field CA1/CA3 pyramidal layer (CA1/CA3-sp), dentate gyrus/granule cell layer (DG-sg), epithalamus (EPI), FT, HPF, hypothalamus (HY), RT, TH, and V3 when mapping to the tissue section (Fig.2E). Using the literature database and the ABMA’s mouse brain tissues^23,24^, a manual annotation was performed to reveal 8 major tissue sub-types. Among them, 7 were consistent with the cluster distribution of the Well-ST-seq-20 μm data (Fig.2F). Through GO analysis, we found that the clusters were predicted to be differentially involved in neural development, signal transduction, and stress response, proving that Well-ST-seq can be used for the mapping of spatial transcriptomics of tissues (Fig.S3C).

**Fig.2.**
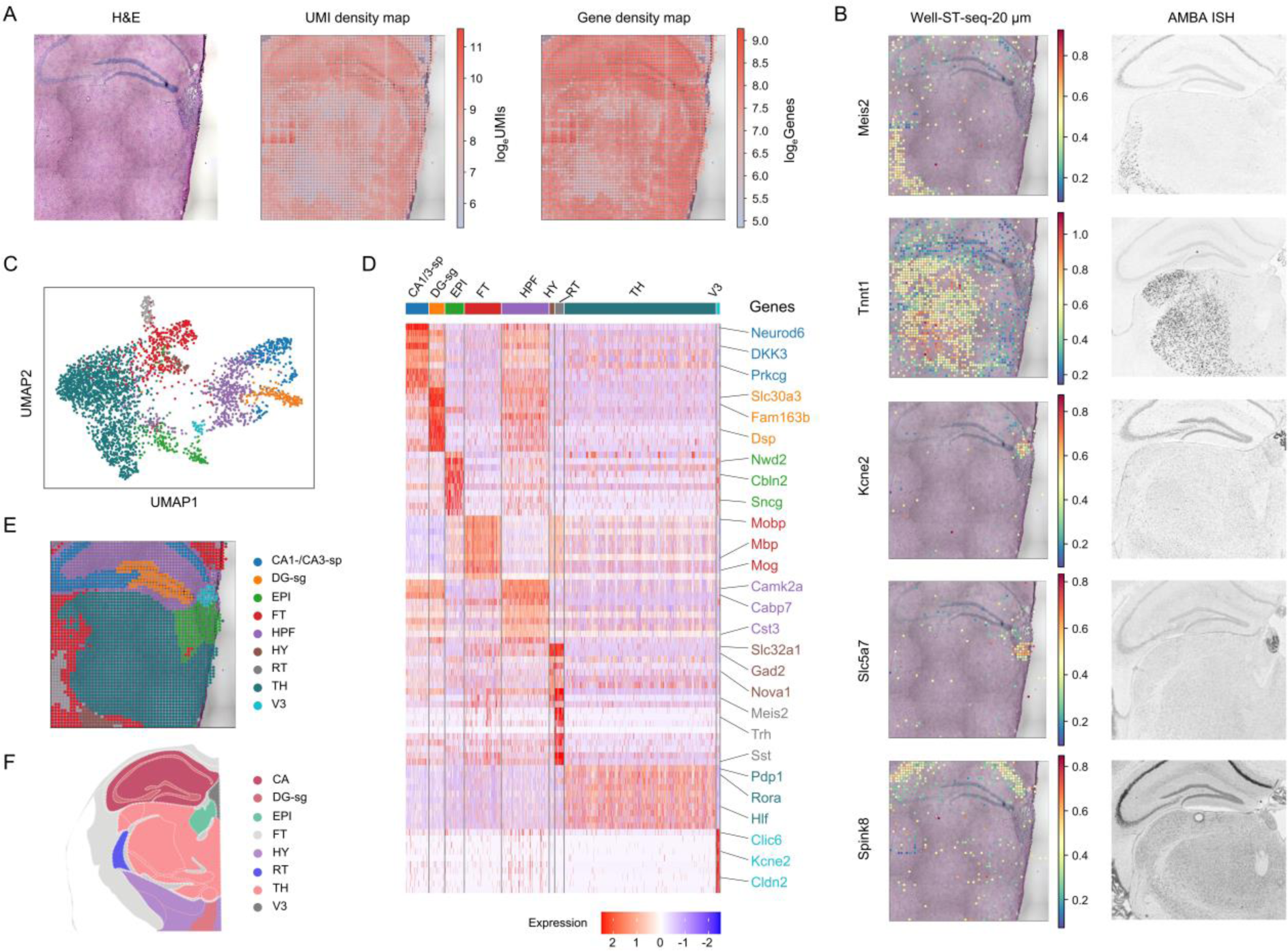
Data analysis of the Well-ST-seq-20 μm. A, H&E staining, UMI and gene density maps of the Well-ST-seq-20 μm. B, Comparison of selected gene expressions detected by Well-ST-seq and the AMBA ISH data. C, Unsupervised clustering analysis shows the clusters of tissue spot transcriptomes. D, Gene expression heatmap of 9 clusters obtained by unsupervised clustering analysis. Top ranked differentially expressed genes are shown in each cluster. E, Overlay of the spatial distribution of the cluster map and tissue image(H&E). F, Anatomic annotation of major tissue regions based on the H&E image and ABMA data.

**Fig.3.**
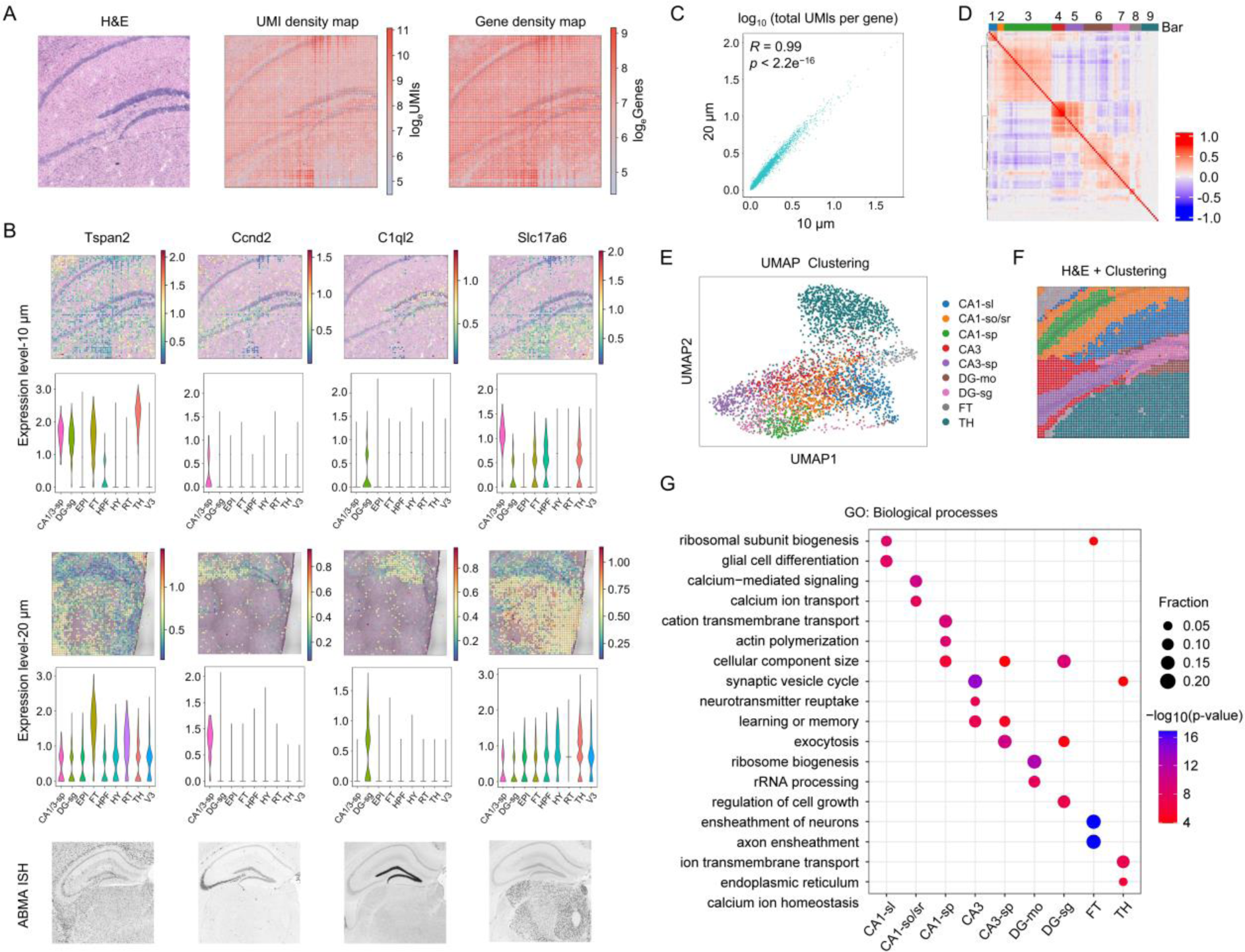
Data analysis of the Well-ST-seq-10 μm. A, H&E staining, UMI and gene density maps of the Well-ST-seq-10 μm. B, Comparison of selected gene expressions detected by Well-ST-seq-10 μm, Well-ST-seq-20 μm, and the AMBA ISH data. C, Reproducibility of Well-ST-seq-10 data between the data from 10 μm- and 20 μm-resolution spatial transcriptomic data in parallel taken as the total UMI counts per gene. Spearman correlation between the total UMI counts per gene of the two datasets is displayed. D, Heatmap showing the normalized expression level of 1,959 genes selected from the disorders genotype-to-phenotype database (DDG2P) in the representative anatomic regions, which showing significant spatial autocorrelation grouped into different clusters. E, Unsupervised clustering analysis shows the clusters of tissue spot transcriptomes. F, Overlay of the spatial distribution of the cluster map and tissue image(H&E). G, Gene Ontology (GO) enrichment analysis of differentially expressed genes in the major clusters in the hippocampus and V. p values, Fisher’s exact test.

### High-spatial-resolution mapping of mouse hippocampus by Well-ST-seq-10 μm

To prove the flexibility of the spatial resolution of Well-ST-seq and conduct high-resolution spatial transcriptomic sequencing of tissue sections, we fabricated 10 μm-resolution microchannels and microwell arrays, and the hippocampus region of the mouse brain was used as the sectioning of tissues (Fig.3A). The UMI and gene heatmaps of the spatial barcode array showed similar distribution when compared to the characteristics of the tissue architecture (Fig.S4A). We also found that the spatial expression mapping and the violin plots of the marker genes in CA, DG, and TH regions were highly consistent between the data of Well-ST-10 μm and Well-ST-20 μm (Fig.3B). The types of up-regulated and down-regulated genes in the two sections were similar (Fig.S4B), and the data of Well-ST-seq also showed highly reproducible between replicates (R = 0.99, Fig.3C). In addition, the expressions of marker genes detected in the data of Well-ST-10 μm and Well-ST-20 μm agreed with the ISH data from the AMBA (Fig.3B). Unsupervised clustering of the spots was performed using their mRNA expression profiles, which showed well-defined spatial patterns, and identified 9 distinct clusters (Fig.3D,E). Compared with the spatial map of the Well-ST-seq-20 μm, Well-ST-seq-10 μm showed a more detailed anatomical morphology corresponding to the HE staining image of the tissue section (Fig.3f), which included FT, CA1, CA3, DG and TH, and the sublayers such as CA1-stratum oriens (CA1-so), CA1-sp, CA1-stratum lacunosum-moleculare (CA1-slm), DG-molecular layer (DG-mo), DG-sg, and CA3-sp. The subsequent clusters matched well with anatomically defined brain regions of the ABMA annotation results, and the marker genes of the clusters were also differentially expressed (Fig.S4C). GO analysis were further performed to identify the biological process of mouse brains and found that it was consistent with the functional enrichment of the hippocampus (Fig.3G)^25^, which proved that the Well-ST-seq technology can be used to mark the region division of fine pathological structures.

### Integration with single-cell sequencing data

Since the resolution of Well-ST-seq can reach 10 μm, which is considered a sweet spot for spatial single-cell analysis^7^, we combined the Well-ST-seq-10 μm data with the reported scRNA-seq data of the mouse brain^26^ to infer the spatial distribution of cell phenotypes in the tissue section. The data of the hippocampus tissue of the scRNA-seq were integrated with Well-ST-seq-10 μm data for unsupervised clustering (Fig.4A), and we found that the spatial data spots matched well with the single-cell transcriptome data. According to the statistical results, ∼ 92% of the genes in the scRNA-seq data were also found in the Well-ST-seq-10 μm data (Fig.4A). The cells in the tissue section contained single and double spots respectively accounted for ∼ 43.7 and ∼ 27.6%, and ∼5100 UMIs on average were detected for per statistical cell, which was comparable to conventional scRNA-seq technology^28^. Then we used single-cell sequencing data as a reference library for the annotation of cell types, allowing the label of cell phenotype directly in each spot, and the visualization of cell distribution in the tissue section can be realized (Fig.4B). Classical neuronal and nonneuronal cell types located in the HPF of the mouse brain, including CA3-pyramidal layer (CA3-Pyr), GABAergic interneuron (GABA), granule cells (GC), astrocyte (Astro), mature oligodendrocytes (MOL), vascular and leptomeningeal cell (VLMC) were mapped to the tissue section, and the landscapes of the cells were consistent with the existing knowledge of the corresponding anatomic regions^22,25,27^ (Fig.4C). Moreover, the endothelial cells and radial glia-like cells (RGL) that should be distributed throughout the hippocampal region were also detected in the tissue section using the inferred results of Well-ST-seq-10 μm (Fig.4C). Finally, to further display the mapping strength of Well-ST-seq for spatial cell annotation in tissue sections, the supervised deconvolution approach was used to estimate the proportional representation of cells, which identified 14 cell types in each simulated spot (Fig.4D), and the cell profiles matched well with the reported cell annotation results, demonstrating that the Well-ST-seq data could be used to both identify functional anatomic regions and cell types for tissue sections. Hence, Well-ST-seq can spatially characterizes the transcriptomic organization and individual cell-type composition of tissues with high resolution and sensitivity.

**Fig.4.**
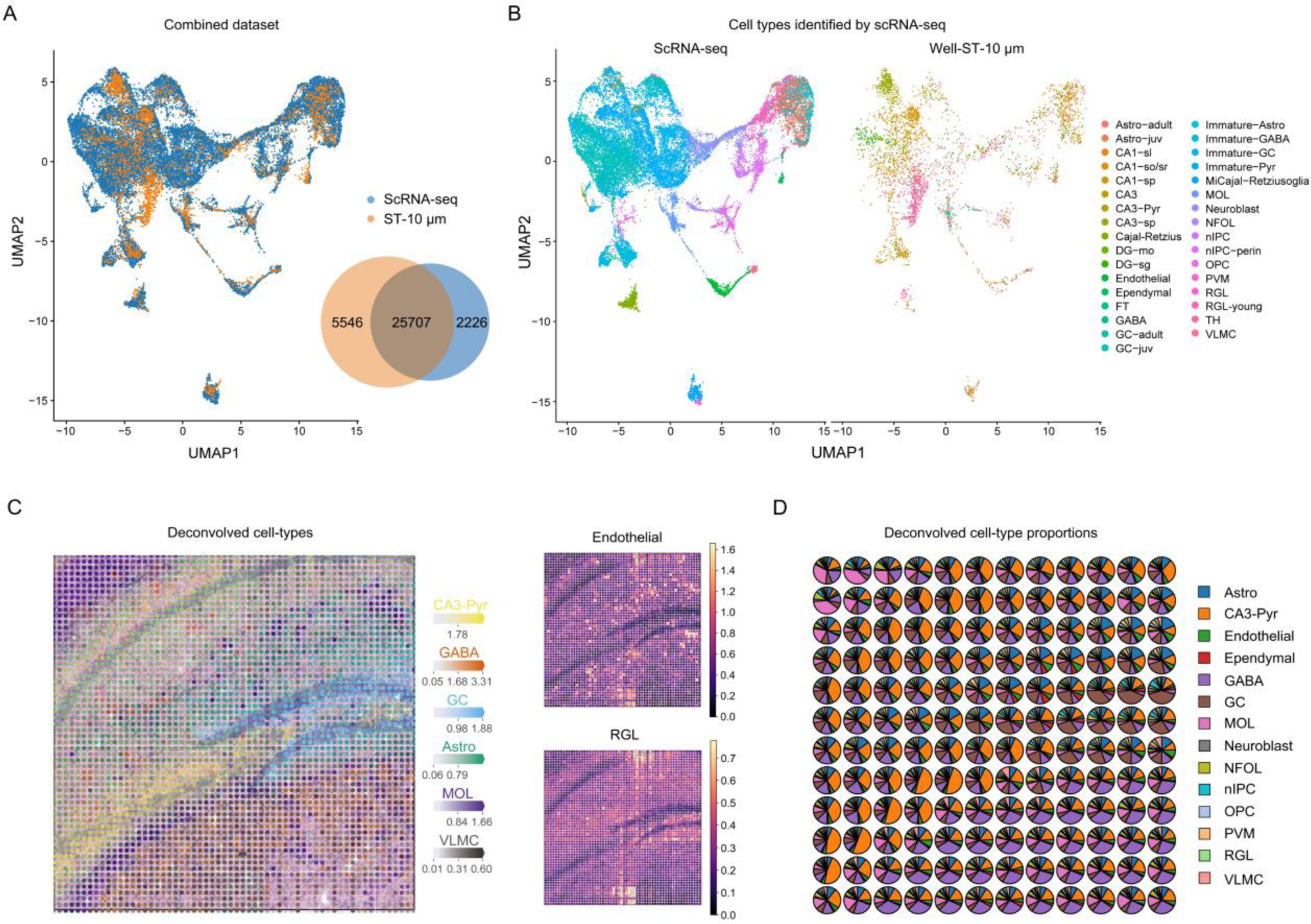
A, Identified cell type profile by Well-ST-seq. A, Integration of scRNA-seq (Hochgerner et al., 2018) and Well-ST-seq-10 μm. The combined data were analyzed with unsupervised clustering. It revealed that the spots of Well-ST-seq-10 μm conformed into the clusters of scRNA-seq data. Inset: venn diagram of genes detected in Well-ST-seq-10 μm data and the scRNA-seq database and their intersections. B, Cell types (different colors) identified by scRNA-seq and comparison with the spots of Well-ST-seq-10 μm. C, Spatial expression pattern of the spots of Well-ST-seq-10 μm in relation to cell types identified (B). D, Spatial distributions of the endothelial cells and RGL in the tissue section. E, Proportions of deconvolved cell types from Well-ST-seq-10 μm for each simulated pixel, including Astro, CA3-Pyr, endothelial, ependymal, GABA, GC, MOL, neuroblast, newly formed oligodendrocyte (NFOL), neuronal intermediate progenitor cells (nIPC), oligodendrocyte precursor cell (OPC), perivascular macrophage (PVM), radial glia-like cells (RGLs), and VLMC.

## Discussion

Based on barcoding beads in microwell arrays, we established a low-cost spatial transcriptomic sequencing technology at cellular level and high RNA capture efficiency. Early spatial sequencing methods are mainly based on multiplexed fluorescent *in-situ* sequencing and hybridization strategies. A major breakthrough in the field arose from the use of spatial barcoding array substrates to *in-situ* capture mRNAs released from a tissue section, and then high throughput NGS sequencing can be used to reconstruct spatial transcriptome maps according to the relationship between the spatial barcodes and the transcripts. Moreover, the measured results of the NGS-based methods are unbiased, genome-wide, and the strategy is more easily adopted by a wider range of scientists in the biological and biomedical research community^13^. Recently, DNA-encoded bead^14,15,22^, HDMI oligonucleotide cluster^16^, and DNA nanoball^17^ were adopted as the RNA-capture substrates for spatial omics sequencing, which improved the spatial resolution of such methods to cellular and subcellular levels. However, these approaches are still demanding, requiring a lengthy and sophisticated step to decode the barcoding sequences. Based on polony-indexed and microfluidic technology, Pixel-seq^18^ and DBiT-seq^19^ have reduced the frequency of the fluorescence decoding period, while the operation proficiency of the methodologies determines the accuracy and repeatability of the technologies. Here, Well-ST-seq can provide a predetermined spatial barcode array at cellular resolution level, which not only eliminates the fluorescence localization step of the barcode array, but also the sequencing process is compatible with the *in-situ* capture of RNA and the NGS technology. Moreover, the transcripts of each annotated cell can reach the level of conventional scRNA-seq on average, which is outstanding among the reported spatial methodologies (Table 1) ^13-19^.

The preparation of Well-ST-seq slide is based on microfluidic technology and does not depend on complex instrument, and the barcode composition of the spot array is deterministic, skipping the fluorescence decoding procedure. The barcoded sequence slide is composed of a reusable PDMS microwell array and magnetic beads, and each microwell array slide consumes only ∼3 g PDMS prepolymer. In addition, the beads with bioactive groups are only filled in the microwell region, which fully reduces the cost for the probe fixation, and 1 μL of the magnetic bead solution (diluted to 15 μL) is sufficient to fill the microwell array larger than 10 mm^2^. Therefore, the consumable cost of Well-ST-seq can reach $1.21/mm^2^ (Table 2), which is much lower than Seq-Scope and Stereo-seq, and is even equivalent to the average cost of the sequenced and stamped gels of Pixel-seq (Table 3). In addition, the preparation cycle of the probe array in Well-ST-seq mainly includes the bead activating and sequence barcoding. The two processes have been optimized and can be performed simultaneously, and the entire barcoding period of the probes in beads can be controlled within 2 h, which is lower than the reported methods (Table 3). Considering the low cost and time consumed of the Well-ST-seq for fabricating the barcoded probe slides, the strategy might open opportunities to break existing limitations of spatial transcriptomics in basic science and clinical work.

Compared with published spatial omics technologies, Well-ST-seq achieved high-sensitivity spatial transcriptomic sequencing (Fig.1F-H). Several factors contributed to the high capture efficiency. First, the cross-linked hydrogel network was coated on the surface of the beads, and the bioactive groups were modified to the chain repeat unit of the hydrogel network, allowing the grafting of high-density RNA-capture probes. Secondly, each microwell spot contains 10-15 beads, and the beads are stacked together in the microwell and exhibit three-dimensional spatial characteristics, which improves the capture area for transcripts. In addition, the spatial characteristics of the microwell and beads composite structures might be beneficial to reduce the molecular crowding between transcripts. Finally, we used microfluidic chips to modify the beads in the microwell with RNA-capture probes, and the continuous flow characteristic of the barcode solution in the microchannel not only keeps the barcode concentration constant, but also improves the reaction efficiency between the bioactive groups, thereby increasing the modification density of the probes.

Another benefit of Well-ST-seq is its scalability and credibility. The slide of Well-ST-seq was prepared by microfluidic chips, and the resolution and probe area for spatial sequencing could be adjusted by changing the arrangement and line width of the microchannels. It is worth mentioning that the bioactive beads are only exist in the microwell, and the non-channel area is blank PDMS surface, which avoids unnecessary modification of probe molecules. Owing to the accurate modification and sweet resolution of its RNA-capture probe array, the spatial distribution of the transcriptomes and cell types analyzed by Well-ST-seq are well matched with the results in AMBA database and reported methods.

In sum, we report on Well-ST-seq technology, providing a low-cost solution for spatial single-cell transcriptome sequencing. Given the demonstrated sensitivity and resolution, Well-ST-seq offers an ideal probe barcoding methodology for the *in-situ* capture of tissue transcripts.

## Limitations of the study

First of all, although the resolution of Well-ST-seq is close to cell level and can be combined with single-cell sequencing data for cell-type analysis, it can only predict cell phenotypes and cannot achieve true single-cell resolution spatial transcriptomes. Secondly, the spatial resolution of Well-ST-seq is limited. The barcode array is prepared by the microfluidic method, and its resolution is limited by the dimension of microchannels, which can theoretically achieve spatial transcriptome sequencing with a resolution of 2 μm. Thirdly, when the number of microchannels is fixed, the resolution of the Well-ST-seq would be inversely proportional to the sequencing area. Of course, the sequencing area can be increased by improving the number of microchannels. Finally, current Well-ST-seq is only for transcriptomes, lacking the display of other omics. In the future, it may be necessary to bind barcode-tagged antibodies and Tn5 transposases to achieve spatial proteomics and chromatin accessibility, respectively.

## Methods

A detailed protocol of the Well-ST-seq procedure is available in the Supplementary Information.

### Animal handling

All relevant procedures involving animal experiments presented in this study are compliant with ethical regulations regarding animal research and were conducted under the approval of by Jilin University Laboratory Animal Center. Food was removed 2 h before the detections were conducted, and the ambient temperature was maintained at 25 °C. Mouse brain was dissected from 12-week-old C57BL/6J female mice. After collection, tissues were snap-frozen in liquid nitrogen prechilled isopentane in Tissue-Tek OCT and transferred to a −80 ℃ freezer for storage before the experiment. Cryosections were cut at a thickness of 10 mm in a Leika CM1950 cryostat. Adult mouse brain was cut coronally.

### Fabrication of the PDMS microchannel and microwell arrays

The PDMS microchannel and microwell arrays were fabricated by soft lithography []. The chrome photomasks with microchannel and microwell patterns were ordered from Shenzhen Nanmei Core Electronics. The molds were fabricated using traditional photolithography and dry etching. Then the degassed PDMS precursor was poured to the molds, and cured at 60 ℃ for 4-6 hours. The solidified PDMS microchannel and microwell were peeled off and punched to inlets and outlets.

### Immobilization of Barcode X

The spatial encoding of the patterned RNA-capture probes on beads was achieved by microfluidic technology. The microfluidic chip contained 60 parallel PDMS microchannels, the microchannel array was aligned on the microwell array (Fig.1C), and a set of barcode X solutions were introduced into the parallel microchannels, respectively. Then the microchannel array was detached from the microwell array. SSC solutions (0.1X) were used to rinse the beads slide.

### Ligation of Barcode Y

After the fixation of the barcode X, the microchannel was replaced by another microchannel to introduce a second set of barcode Y along the orthogonal direction of the barcode X fluids. After the extension and denaturation of the barcode X and barcode Y, spatially barcoded RNA-capture sequences were respectively fixed on the beads in the microwell array.

### Data processing

To ensure the data’s veracity and integrity, and subject it to a meticulous quality control process, genes exhibiting cell counts below the predetermined threshold of 10 were eliminated via the filter_genes function in Scanpy(2). The remaining genes were subsequently normalized using the normalize_total function, and to mitigate any skewed distribution, the normalized values underwent a log transformation through the log1p function.

### Benchmark

To compare the similarities and differences between our sequencing technology and other sequencing technologies, we compared the similarities and differences of the sum of the expression of all genes in per cell (nCount_RNA) between the spatial transcriptome 10um data and four external data: ST_DBi_10um(), ST_10X_55um(), ST_scope_0.8um(), ST_Stereo_0.7um().

### Data integration and clustering

To discern the intricate nuances of gene expression in a spatially variant manner, SpaGCN(2) was employed to amalgamate spatial transcriptomics expression matrices, spatial location, and histological images information, and to cluster each spot based on the integrated information. During the clustering process, in addition to the default parameters, parameters where (l=0.5 res=20, lr=0.05, max_epochs=200) was used in the spatial transcriptome data set of 10 microns to obtain 13 clusters, rather than 29 clusters obtained in the data set of 20microns using parameters (l=0.9 res=20, lr=0.05, max_epochs=200).

### Differential expression analysis and annotation

In order to discern the differentially expressed genes (DEGs) within each cluster, differential expression analysis was performed using Scanpy. And the rank_genes_groups function was utilized with the default parameters to identify cluster-specific biomarkers, which allowed for an accurate and comprehensive analysis of the gene expression data.

After the annotation of the cell type by observing the expression of the recognised markers, 9 major cell types was obtained for 10-microns-data including CA1-sl(glul), CA1-so/sr(git1), CA1-sp(hpca), CA3(cnr1), CA3-sp(snca), DG-mo(atp1a2), DG-sg(dsp), FT(cldn11), TH(prkcd). Another 9 major cell types was also annotated for 20-microns-data including CA1-sp/CA3-sp(neurod6), DG-sg(2010300c02rik), EPI(calb2), FT(mobp), HPF(ddn), HY(slc32a1), TH(pdp1), RT(six3), V3(clic6).

### Integration of single-cell data and spatial transcriptome data

In order to demonstrate the similarity between scRNA-seq (GSE104323, https://www.ncbi.nlm.nih.gov/geo/query/acc.cgi?acc=GSE104323) data and Spatial RNA-seq data, and to provide a theoretical foundation for the subsequent process of cell type mapping, we integrated spatial transcriptome data into single-cell data utilizing Seurat(3) Integration algorithm. To ensure the veracity of the data, we employed the SCT_transform normalization method and meticulously selected the top 3000 most variable genes in the single-cell data during the Seurat Integration process. We utilized the Seurat package to perform various analyses, such as dimensionality reduction, clustering, and identification of differentially expressed genes. To perform dimensionality reduction, we initially computed the PCA matrix based on the integrated data using the RunPCA function in the Seurat standard workflow. Subsequently, we constructed a shared nearest neighbor graph between every cell and its k nearest neighbors by calculating the k nearest neighbors using the FindNeighbors function with 30 components. This nearest neighbor graph was used to find clusters by leveraging the Louvain algorithm (4) with 0.1 resolution via the FindClusters function.To visualize the clustering results, we utilized the Uniform Manifold Approximation and Projection (UMAP) method via the RunUMAP function with 30 components. Overall, these sophisticated methods allowed us to accurately integrate spatial transcriptome data into single-cell data, and the subsequent analyses provided valuable insights into the intricacies of the gene expression patterns.

In order to annotate the cell type of each spot in spatial transcriptome data more finely, or to distinguish the proportional state of the cell type in each spot, cell2location(5), a sophisticated algorithm, was used to seamlessly integrate single-cell data into spatial transcriptome data. By leveraging this algorithm, the probability of the cell type from single-cell data was inferred in each spot. And by judiciously comparing the probabilities, the major cell type in each spot was precisely deduced.

### Gene Ontology enrichment analysis

We defined specific differential expressed genes (DEGs) in different groups of each cell type. Based on these genes, enriched gene ontology (GO) terms were then acquired for each cluster or group using R package clusterProfiler(v4.2.2) (6) with default parameters. Annotation Database org.Mm.eg.db was preformed to map these DEGs, and the visualization was showed by bar plot.

### Statistical analysis

To identified differentially expressed genes between two groups of cells, we used a t-test. We also used Benjamini Hochberg algorithm to correct the p-values to identify the GO terms.

## Data and code availability

The accession number for the sequencing data reported in this paper is submitted to GEO:

## Acknowledgements

This work was supported by the National Natural Science Foundation of China (Grant no. 22275071), and the China Postdoctoral Science Foundation (2020M681047).

## Author contributions

## Competing interests

The authors declare that they have no known competing financial interests or personal relationships that could have appeared to influence the work reported in this paper.

## Additional information

**Extended data** are available for this paper at.

**Supplementary Information** The online version contains supplementary material available at.

**Correspondence and requests for materials** should be addressed to Junhu Zhang or Chongyang Liang.

## Material availability

All materials used for Well-ST-seq are commercially available.

## SUPPLEMENTARY INFORMATION

**Fig.S1.**
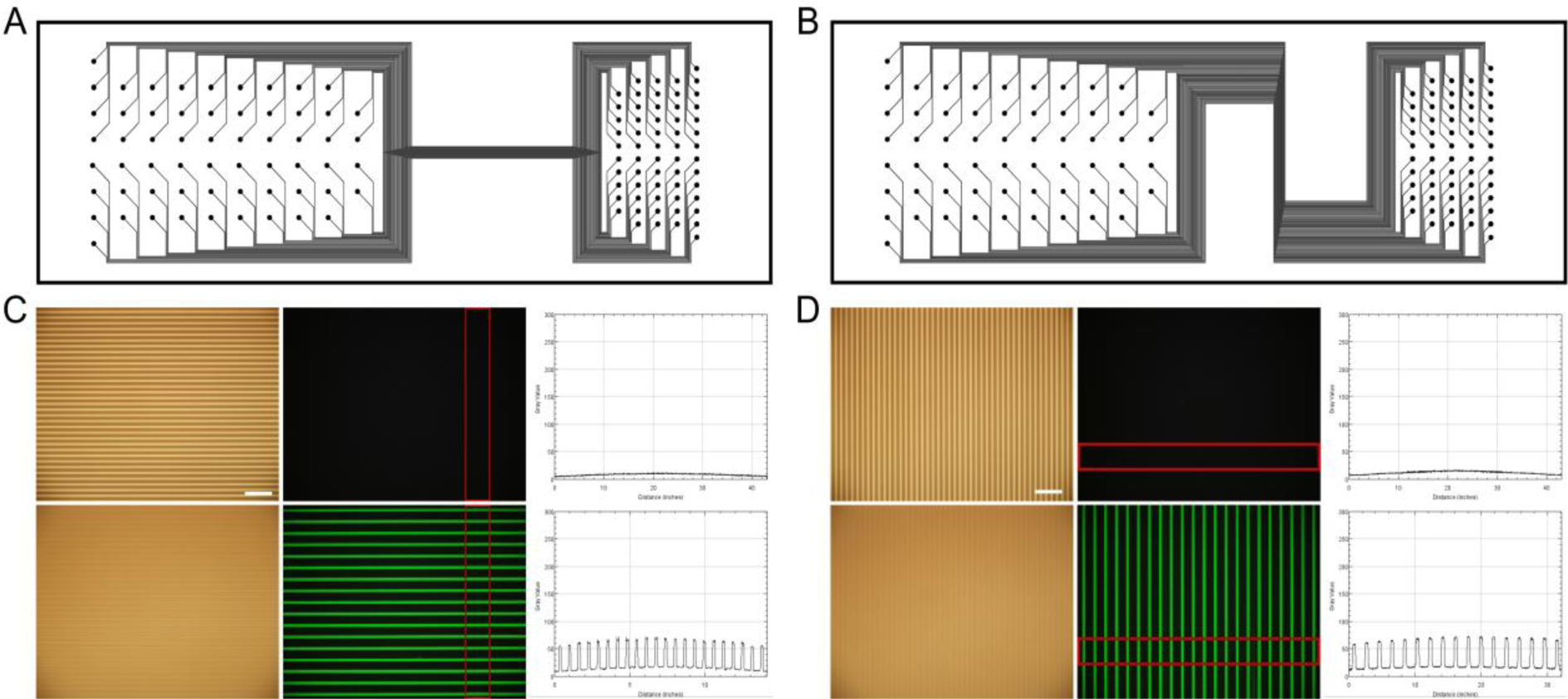
Microchannel design of Well-ST-Seq and Validation. A-B, AutoCAD design of the PDMS chips with 10 μm channel width. C,D, Evaluation of the microchannel flowability and independence (10 μm microchannel width). (Upper) Optical, fluorescence microscope images, and line profile of the fluorescence intensity of the microchannels without liquid infusion. (Lower) Optical, fluorescence microscope images, and line profile of the fluorescence intensity of the fluorophore-FAM (Green) labeled barcode solution and water that were alternately introduced into the microchannels. The optical microscope images show that the liquids can flow through the microchannels, and the fluorescence microscope images show that the fluorescence signal only appears in the microchannels that flow into the labeled barcode solutions, which proves that there is no crosstalk between the neighboring microchannels. Scale bar = 100 μm.

**Fig.S2.**
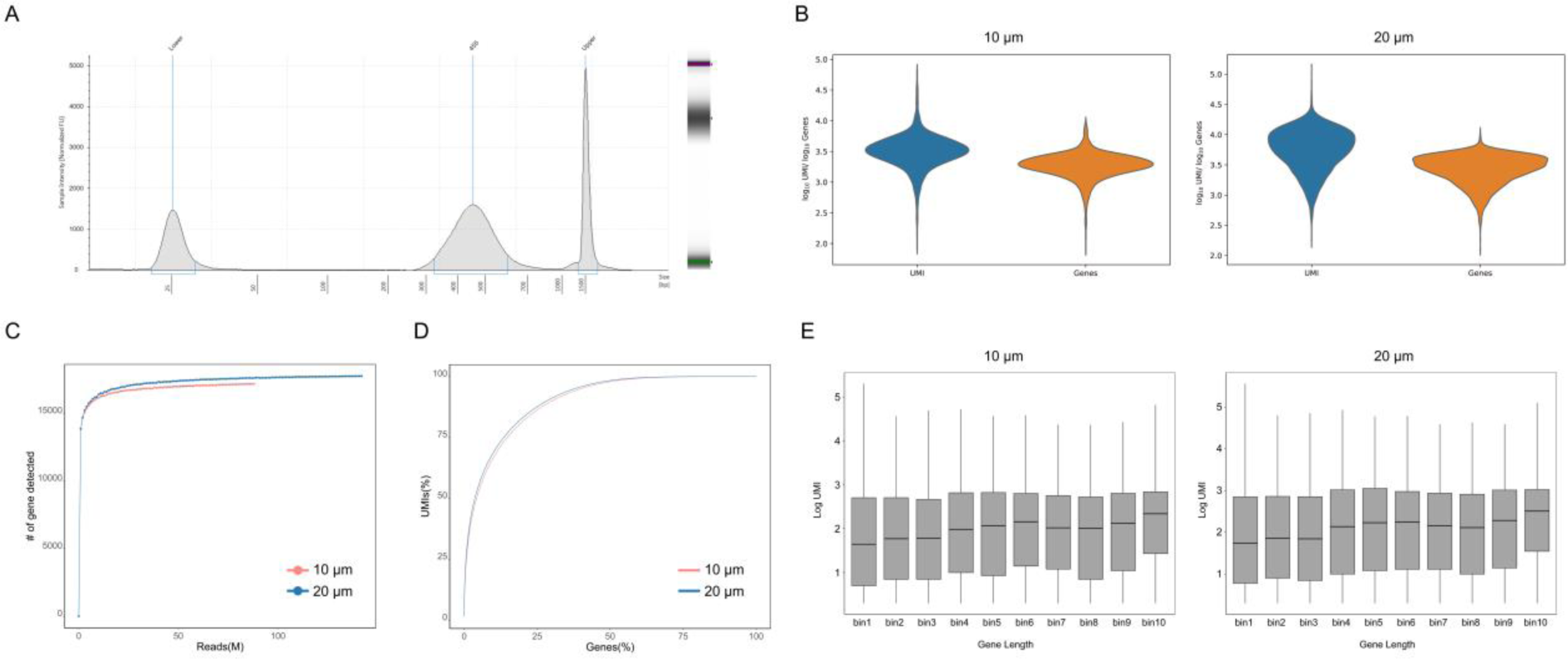
Data quality evaluation. A, Size distribution of library sequences form Well-ST-seq-10 μm. B, Violin plots showing the number of detected UMI and genes of Well-ST-seq-10 μm and Well-ST-seq-20 μm. C, The saturation curves of the data of Well-ST-seq-10 μm and Well-ST-seq-20 μm. D, Cumulative number of genes versus UMI. Top 10% genes account for 70% of the UMIs and top 30% genes account for 85% of UMIs. E, Gene length bias analysis of Well-ST-seq, and shorter genes having lower UMI counts.

**Fig.S3.**
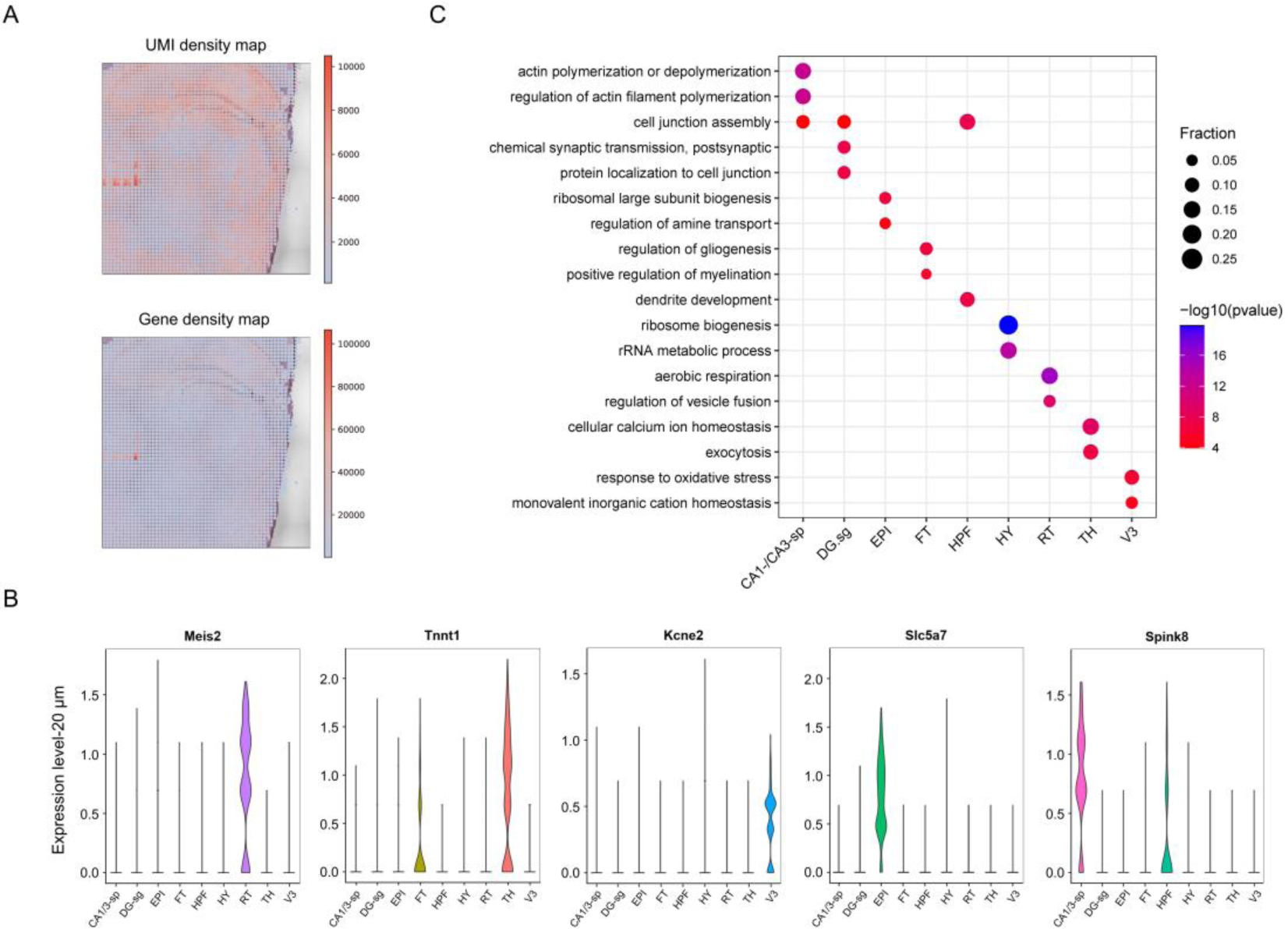
Data analysis of the Well-ST-seq-20 μm. A, Overlay of the spatial distribution of the UMI, gene density map, and tissue image (H&E). B, Violin plots of selected top genes showing differential expression in each cluster. C, GO enrichment analysis of differentially expressed genes in the major clusters in the mouse brain and V. p values, Fisher’s exact test.

**Fig.S4.**
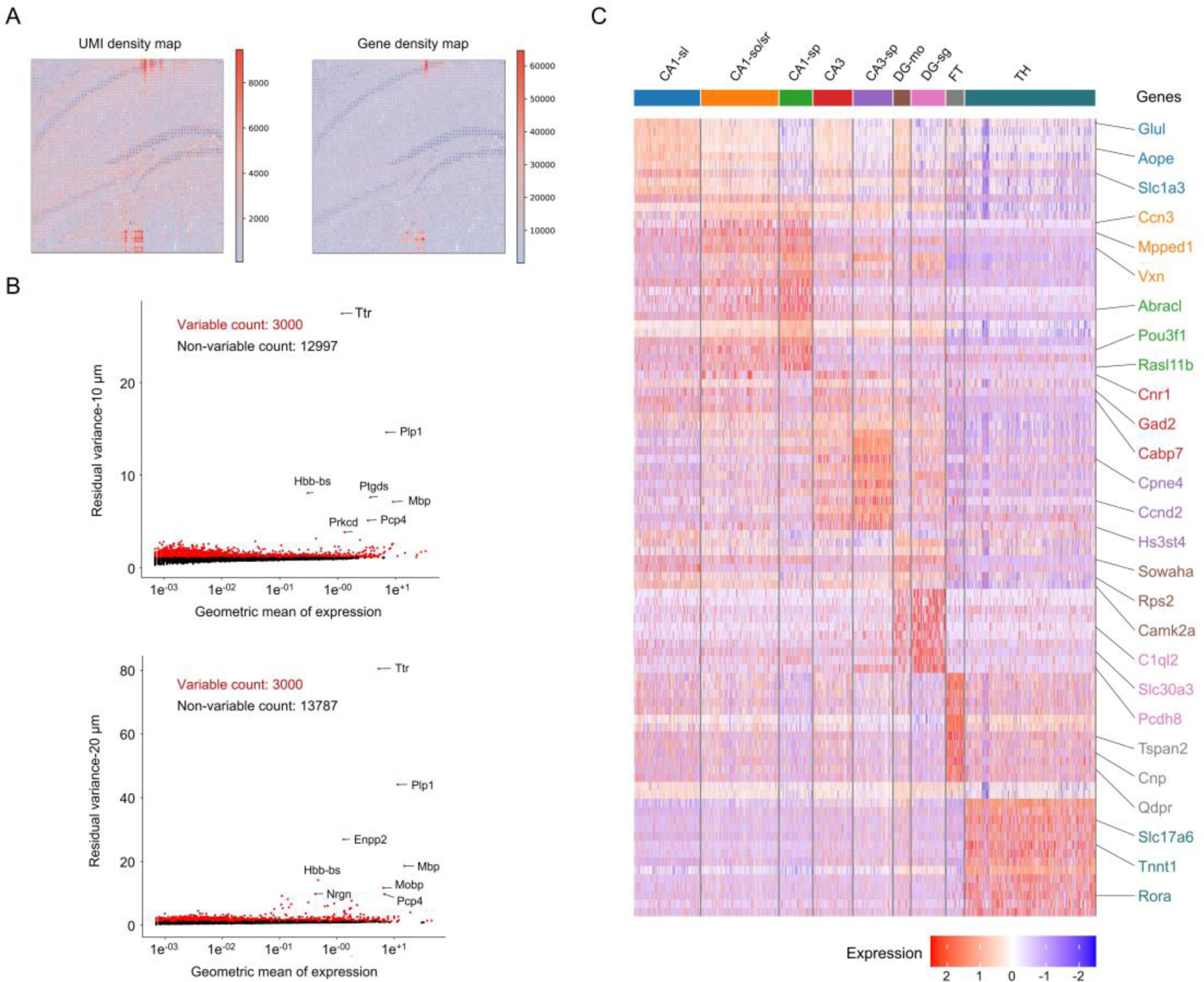
Data analysis of the Well-ST-seq-10 μm. A, Overlay of the spatial distribution of the UMI, gene density map, and tissue image (H&E). B, The up-regulated and down-regulated genes of the data of Well-ST-seq-10 μm and Well-ST-seq-20 μm. C, Gene expression heatmap of 9 clusters obtained by unsupervised clustering analysis. Top ranked differentially expressed genes are shown in each cluster.

**Table S1.**
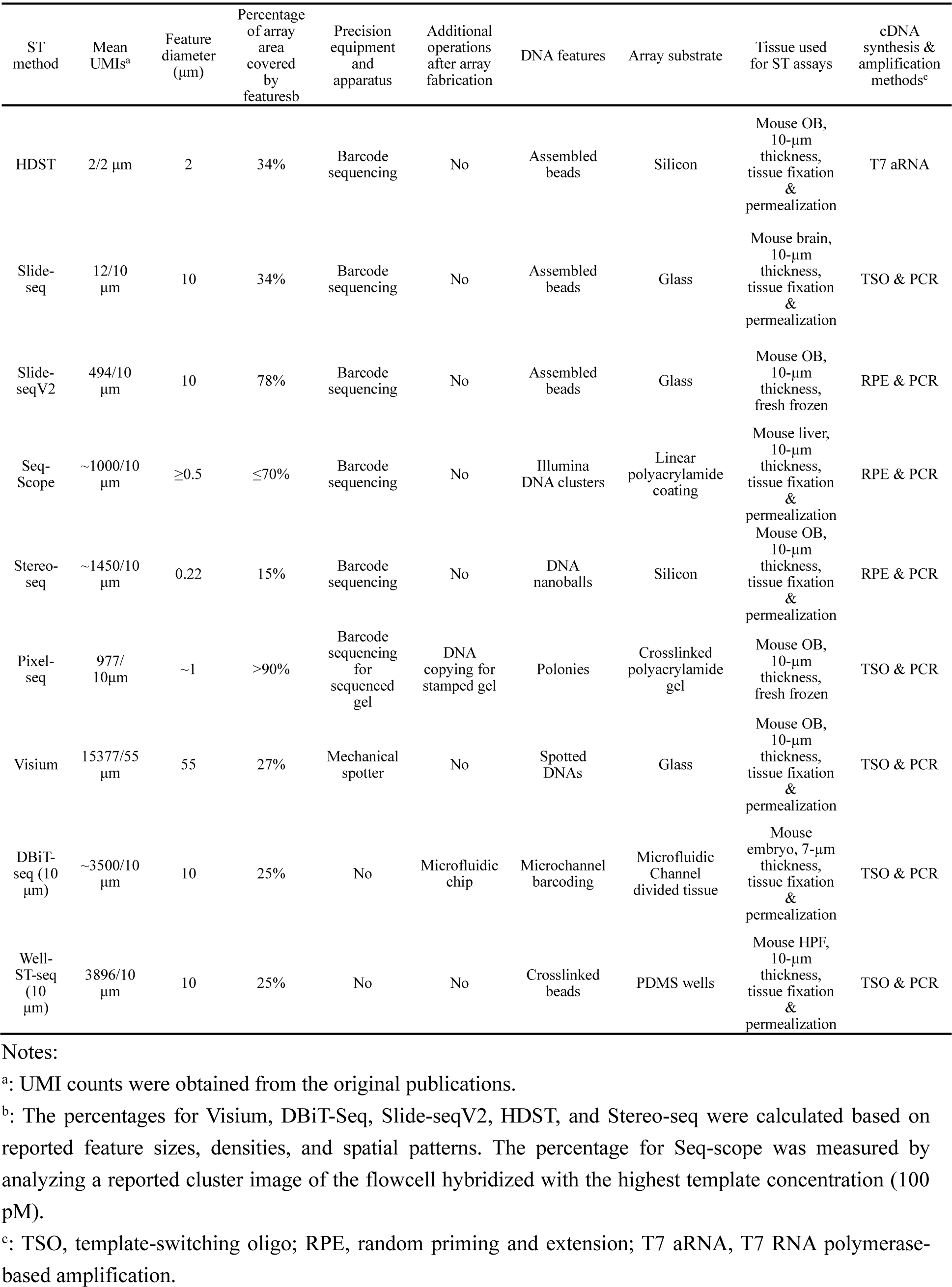
Comparison of DNA array-based spatial transcriptomic assays.

| ST method | Mean UMIs<sup>a</sup> | Feature diameter (μm) | Percentage of array area covered by features<sup>b</sup> | Precision equipment and apparatus | Additional operations after array fabrication | DNA features | Array substrate | Tissue used for ST assays | cDNA synthesis & amplification methods<sup>c</sup> |
| --- | --- | --- | --- | --- | --- | --- | --- | --- | --- |
| HDST | 2/2 μm | 2 | 34% | Barcode sequencing | No | Assembled beads | Silicon | Mouse OB, 10-μm thickness, tissue fixation & permeabilization | T7 aRNA |
| Slide-seq | 12/10 μm | 10 | 34% | Barcode sequencing | No | Assembled beads | Glass | Mouse brain, 10-μm thickness, tissue fixation & permeabilization | TSO & PCR |
| Slide-seqV2 | 494/10 μm | 10 | 78% | Barcode sequencing | No | Assembled beads | Glass | Mouse OB, 10-μm thickness, fresh frozen | RPE & PCR |
| Seq-Scope | ~1000/10 μm | ≥0.5 | ≤70% | Barcode sequencing | No | Illumina DNA clusters | Linear polyacrylamide coating | Mouse liver, 10-μm thickness, tissue fixation & permeabilization | RPE & PCR |
| Stereo-seq | ~1450/10 μm | 0.22 | 15% | Barcode sequencing | No | DNA nanoballs | Silicon | Mouse OB, 10-μm thickness, tissue fixation & permeabilization | RPE & PCR |
| Pixel-seq | 977/10 μm | ~1 | >90% | Barcode sequencing for sequenced gel | DNA copying for stamped gel | Polonies | Crosslinked polyacrylamide gel | Mouse OB, 10-μm thickness, fresh frozen | TSO & PCR |
| Visium | 15377/55 μm | 55 | 27% | Mechanical spotter | No | Spotted DNAs | Glass | Mouse OB, 10-μm thickness, tissue fixation & permeabilization | TSO & PCR |
| DBiT-seq (10 μm) | ~3500/10 μm | 10 | 25% | No | Microfluidic chip | Microchannel barcoding | Microfluidic Channel divided tissue | Mouse embryo, 7-μm thickness, tissue fixation & permeabilization | TSO & PCR |
| Well-ST-seq (10 μm) | 3896/10 μm | 10 | 25% | No | No | Crosslinked beads | PDMS wells | Mouse HPF, 10-μm thickness, tissue fixation & permeabilization | TSO & PCR |

**Notes:**
<sup>a</sup>: UMI counts were obtained from the original publications.
<sup>b</sup>: The percentages for Visium, DBiT-Seq, Slide-seqV2, HDST, and Stereo-seq were calculated based on reported feature sizes, densities, and spatial patterns. The percentage for Seq-scope was measured by analyzing a reported cluster image of the flowcell hybridized with the highest template concentration (100 pM).
<sup>c</sup>: TSO, template-switching oligo; RPE, random priming and extension; T7 aRNA, T7 RNA polymerase-based amplification.

**Table S2.**
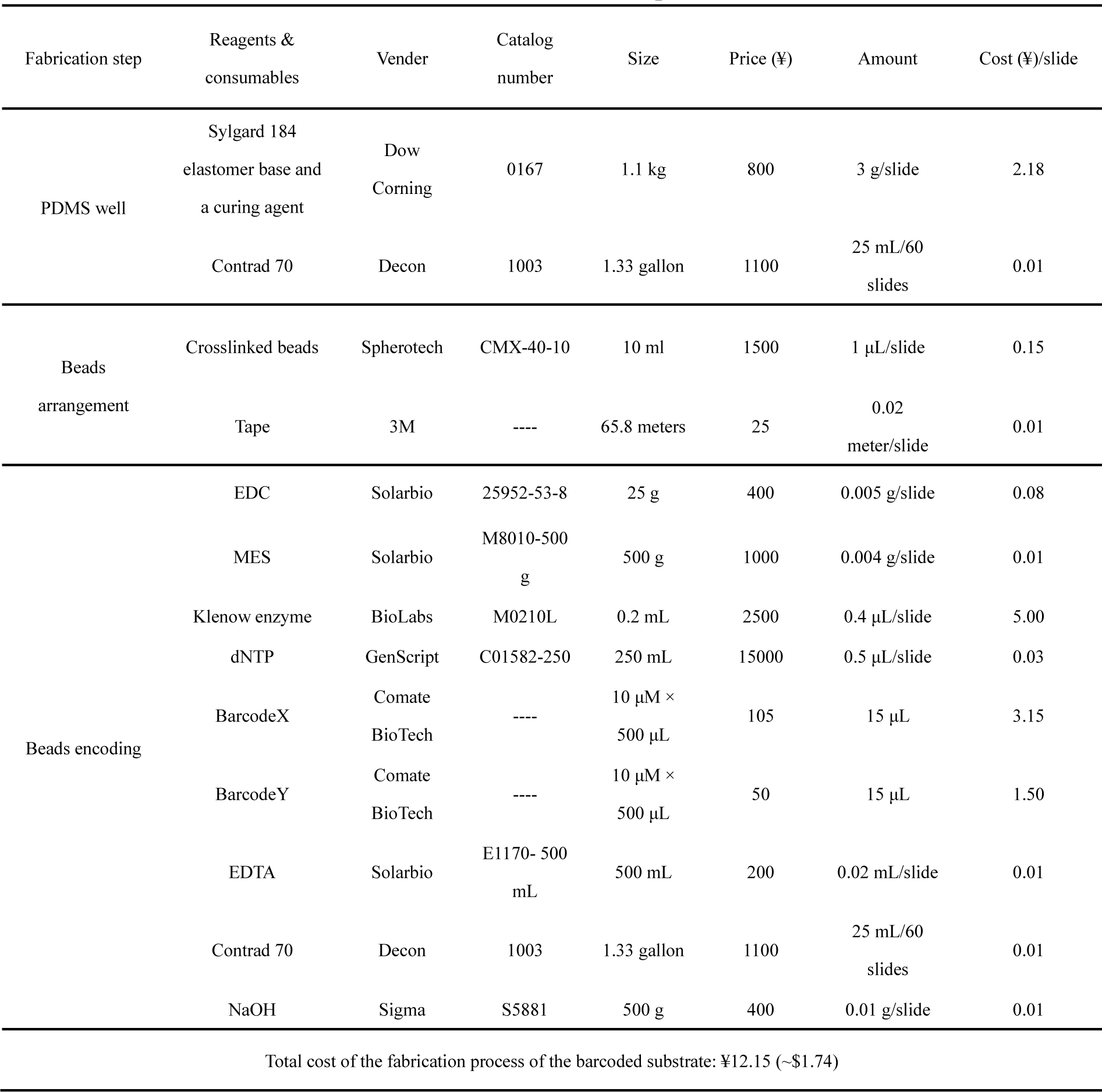
Consumable lists for the fabrication of Well-ST-seq slide.

**Table S3.**
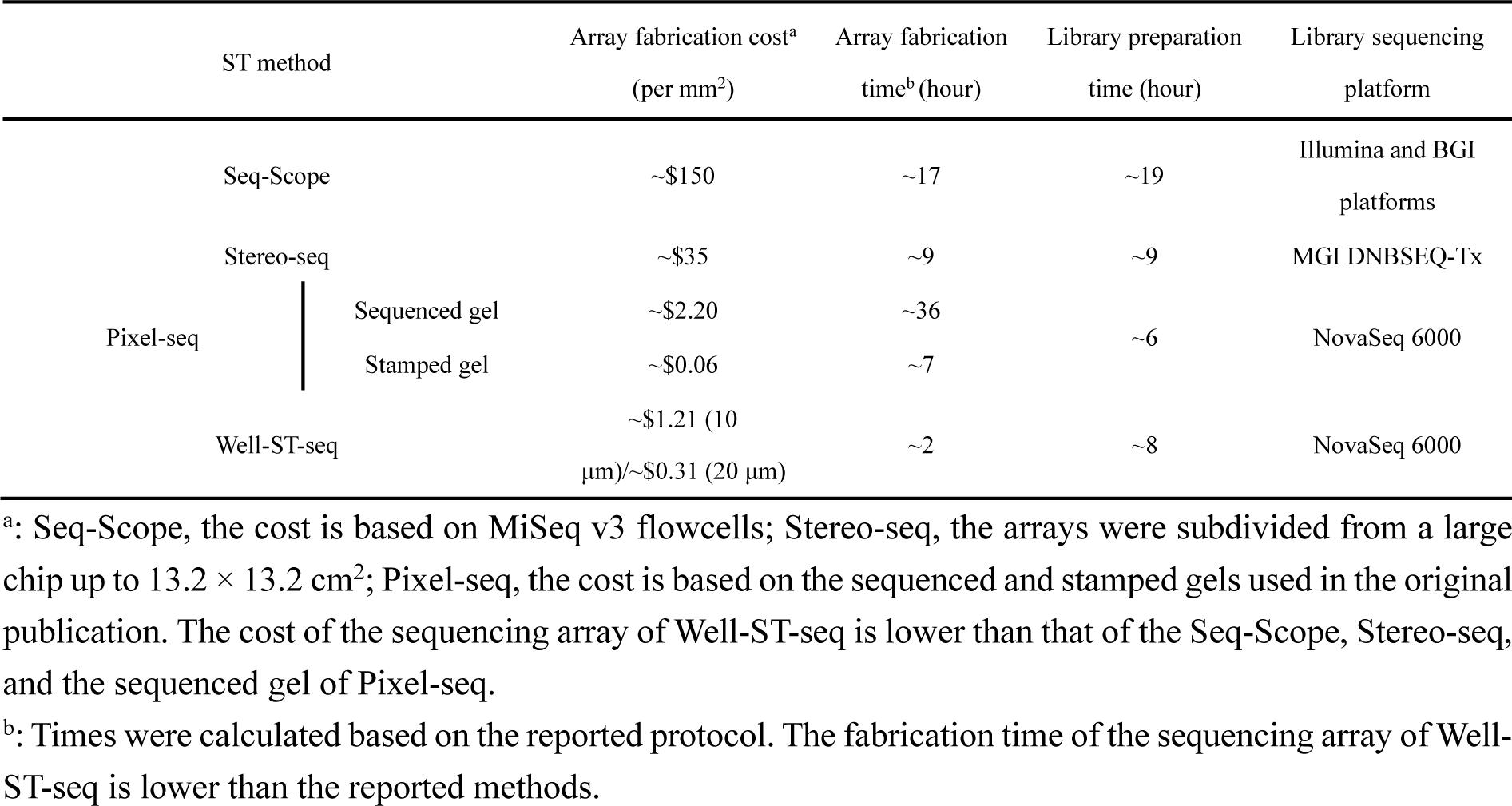
Comparison of Seq-Scope, Stereo-seq, and Pixel-seq assays.

## Notes

### Competing Interest Statement

The authors have declared no competing interest.

## References

1. A. Rao, D. Barkley, G. S. Franca, I. Yanai, Nature 2021, 596, 211.

2. S. M. Lewis, M.-L. Asselin-Labat, N. Quan, J. Berthelet, X. Tan, V. C. Wimmer, D. Merino, K. L. Rogers, S. H. Naik, Nature Methods 2021, 18, 997.

3. S. K. Longo, M. G. Guo, A. L. Ji, P. A. Khavari, Nature Reviews Genetics 2021, 22, 627.

4. L. Zhang, D. Chen, D. Song, X. Liu, Y. Zhang, X. Xu, X. Wang, Signal Transduction and Targeted Therapy 2022, 7.

5. M. Eisenstein, Nature 2022, 601, 658.

6. L. Moses, L. Pachter, Nature Methods 2022, 19, 534.

7. V. Marx, Nature Methods 2021, 18, 9.

8. J. M. Levsky, S. M. Shenoy, R. C. Pezo, R. H. Singer, Science 2002, 297, 836.

9. J. H. Lee, E. R. Daugharthy, J. Scheiman, R. Kalhor, J. L. Yang, T. C. Ferrante, R. Terry, S. S. F. Jeanty, C. Li, R. Amamoto, D. T. Peters, B. M. Turczyk, A. H. Marblestone, S. A. Inverso, A. Bernard, P. Mali, X. Rios, J. Aach, G. M. Church, Science 2014, 343, 1360.

10. C. Xia, J. Fan, G. Emanuel, J. Hao, X. Zhuang, Proceedings of the National Academy of Sciences of the United States of America 2019, 116, 19490.

11. C.-H. L. Eng, M. Lawson, Q. Zhu, R. Dries, N. Koulena, Y. Takei, J. Yun, C. Cronin, C. Karp, G.-C. Yuan, L. Cai, Nature 2019, 568, 235.

12. L. Larsson, J. Frisen, J. Lundeberg, Nature Methods 2021, 18, 15.

13. P. L. Stahl, F. Salmen, S. Vickovic, A. Lundmark, J. F. Navarro, J. Magnusson, S. Giacomello, M. Asp, J. O. Westholm, M. Huss, A. Mollbrink, S. Linnarsson, S. Codeluppi, A. Borg, F. Ponten, P. I. Costea, P. Sahlen, J. Mulder, O. Bergmann, J. Lundeberg, J. Frisen, Science 2016, 353, 78.

14. S. G. Rodriques, R. R. Stickels, A. Goeva, C. A. Martin, E. Murray, C. R. Vanderburg, J. Welch, L. M. Chen, F. Chen, E. Z. Macosko, Science 2019, 363, 1463.

15. S. Vickovic, G. Eraslan, F. Salmen, J. Klughammer, L. Stenbeck, D. Schapiro, T. Aijo, R. Bonneau, L. Bergenstrahle, J. F. Navarro, J. Gould, G. K. Griffin, A. Borg, M. Ronaghi, J. Frisen, J. Lundeberg, A. Regev, P. L. Stahl, Nature Methods 2019, 16, 987.

16. C.-S. Cho, J. Xi, Y. Si, S.-R. Park, J.-E. Hsu, M. Kim, G. Jun, H. M. Kang, J. H. Lee, Cell 2021, 184, 3559.

17. A. Chen, S. Liao, M. Cheng, K. Ma, L. Wu, Y. Lai, X. Qiu, J. Yang, J. Xu, S. Hao, X. Wang, H. Lu, X. Chen, X. Liu, X. Huang, Z. Li, Y. Hong, Y. Jiang, J. Peng, S. Liu, M. Shen, Cell 2022, 185, 1777.

18. X. Fu, L. Sun, R. Dong, J. Y. Chen, R. Silakit, L. F. Condon, Y. Lin, S. Lin, R. D. Palmiter, L. Gu, Cell 2022, 185, 4621.

19. Y. Liu, M. Yang, Y. Deng, G. Su, A. Enninful, C. C. Guo, T. Tebaldi, D. Zhang, D. Kim, Z. Bai, E. Norris, A. Pan, J. Li, Y. Xiao, S. Halene, R. Fan, Cell 2020, 183, 1665.

20. K. Vandereyken, A. Sifrim, B. Thienpont, T. Voet, Nature Reviews Genetics 2023, DOI: 10.1038/s41576-023-00580-2.

21. J. F. Navarro, J. Sjostrand, F. Salmen, J. Lundeberg, P. L. Stahl, Bioinformatics 2017, 33, 2591.

22. R. R. Stickels, E. Murray, P. Kumar, J. Li, J. L. Marshall, D. J. Di Bella, P. Arlotta, E. Z. Macosko, F. Chen, Nature Biotechnology 2021, 39, 313.

23. W.-T. Chen, A. Lu, K. Craessaerts, B. Pavie, C. S. Frigerio, N. Corthout, X. Qian, J. Lalakova, M. Kuhnemund, I. Voytyuk, L. Wolfs, R. Mancuso, E. Salta, S. Balusu, A. Snellinx, S. Munck, A. Jurek, J. F. Navarro, T. C. Saido, I. Huitinga, J. Lundeberg, M. Fiers, B. De Strooper, Cell 2020, 182, 976.

24. X. Qian, K. D. Harris, T. Hauling, D. Nicoloutsopoulos, A. B. Munoz-Manchado, N. Skene, J. Hjerling-Leffler, M. Nilsson, Nature Methods 2020, 17, 101.

25. E. S. Lein, M. J. Hawrylycz, N. Ao, M. Ayres, A. Bensinger, A. Bernard, A. F. Boe, M. S. Boguski, K. S. Brockway, E. J. Byrnes, L. Chen, L. Chen, T.-M. Chen, M. C. Chin, J. Chong, B. E. Crook, A. Czaplinska, Nature 2007, 445, 168.

26. H. Hochgerner, A. Zeisel, P. Lobnnerberg, S. Linnarsson, Nature Neuroscience 2018, 21, 290.

27. A. Saunders, E. Z. Macosko, A. Wysoker, M. Goldman, F. M. Krienen, H. de Rivera, E. Bien, M. Baum, L. Bortolin, S. Wang, A. Goeva, J. Nemesh, N. Kamitaki, S. Brumbaugh, D. Kulp, S. A. McCarroll, Cell 2018, 174, 1015.

28. J. Ding, X. Adiconis, S. K. Simmons, M. S. Kowalczyk, C. C. Hession, N. D. Marjanovic, T. K. Hughes, M. H. Wadsworth, T. Burks, L. T. Nguyen, J. Y. H. Kwon, B. Baraks, W. Ge, A. J. Kedaigle, S. Carroll, S. Li, N. Hacohen, O. Rozenblatt-Rosen, A. K. Shalek, A.-C. Villani, A. Regev, J. Z. Levin, Nature Biotechnology 2020, 38, 737.

